# Multiscale Correlations between Joint and Tissue-Specific Biomechanics and Anatomy in Postmortem Ovine Stifles

**DOI:** 10.1101/2024.03.15.585312

**Authors:** Aritra Chatterjee, Zachary Robert Davis, Timothy Lescun, Deva D. Chan

## Abstract

The stability of the knee joint is an important indicator of its overall health and function. Joint stability depends on multiple structural and functional properties that include the anatomy of the underlying bones, the geometry and stiffness of the joint capsule and the soft tissues within like tendon, ligaments, cartilage and meniscus. The multiscale biomechanical relationships between the form and function of the whole joint and individual tissues can provide useful insights on the physiological condition of the knee and require further exploration. To better understand these relationships, in this study we compare multiple structural and mechanical parameters in healthy ovine stifles (n = 6). Specifically, we have evaluated joint laxity, joint morphology, individual tissue *T_2_\** relaxation and mechanical properties of the anterior and posterior cruciate ligaments (ACL, PCL), medial and lateral collateral ligaments (MCL, LCL), the patellar tendon, menisci, and cartilage from the femoral condyles. Using mechanical testing at two length scales along with anatomical and quantitative magnetic resonance imaging (MRI) scans, we investigate the correlation between individual tissue and whole joint mechanical properties. We also performed statistical tests to measure the strength of correlation coefficients between the measured metrics at multiple levels among whole joint mechanics, joint size, and individual tissue properties. We observe positive correlations between the joint laxity forces and the epicondyle-to-epicondyle distance measured as an anatomical marker. We also find that the viscoelastic properties of the tendons and ligaments correlate positively with joint laxity forces. No such correlations were observed between the cartilage and meniscus properties and the joint laxity forces. Further, we found a weak inverse correlation between the tissue viscoelastic properties and *T_2_\** for MCL; strong and moderately positive correlations for cartilage samples from both femoral condyles and the menisci, LCL and PCL respectively. These results provide useful insights into the differential role of individual tissue properties that can be used to predict the whole joint responses that are key indicators of knee health and performance.

## INTRODUCTION

The knee is subject to a wide range of complex loading conditions that depend on multiple factors such as the body weight of the individual, the frequency and extent of movement of the limbs, and their energy absorption capacity [1]. Despite its importance in everyday mobility and quality of life, the knee is susceptible to different types of injuries, affecting nearly 46% adults over their lifetime [2]. Injuries to or impairment of the knee can significantly impact human mobility and well-being [3]. The anatomic structures within the knee, the connective tissues including the tendons, ligaments together with the meniscus and articular cartilage all contribute significantly influencing the movement and stability of the joint [4]. A thorough understanding of the structure and functional properties of the knee joint, and its underlying structures are required to identify the cause of such injuries and develop better diagnostic or precautionary measures [5]. Multiscale biomechanical studies of the knee joint in large animal models that can closely capture the human joint anatomy and loading conditions, are often useful in designing studies. Owing to their low-cost relative to other large animals, and physiological similarities with humans, ovine models are commonly used in orthopedic research and late-stage preclinical studies [6].

Joint laxity, often widely used as a metric to quantify the stability of the joint, is defined as a measurement of the net joint movement under the application of an external force during a state of muscular relaxation [7]. Quantification of joint laxity is of clinical importance, as it is linked to joint instability which can arise because of injuries to soft tissues like ligament tear or in case of degenerative diseases like osteoarthritis [8]. Anterior drawer tests are a routinely used technique to quantify joint laxity in a clinical setting. [9]. Laxity depends on multiple factors including the mechanical constraints of the underlying soft tissues, tibio-femoral bone shape and multiple other factors [10]. Investigating the specific correlations between the individual tissue mechanics and structure with joint laxity can help us understand the relationship between the macroscale joint mechanics and the aggregate mechanical properties of the soft tissues.

Magnetic resonance imaging (MRI) is a widely used technique to image knee anatomy [11], and quantitative relaxometry is sensitive to tissue composition including changes in water content, proteoglycans, and collagen fiber orientation [12]. Both transverse relaxation time (*T_2_*) and effective *T_2_* (*T_2_\**) have been shown to correlate to cartilage degeneration [13]. An increase in the *T_2_* correlates with damage to articular cartilage, signifying deterioration of the collagen network and increase in the water content [14]. *T_2_\** has also correlated with collagen fiber orientation, water content, and tissue stiffness [15, 16]. In this study, we use *T_2_\** mapping to determine how well overall joint and individual mechanics correlate with changes in *T_2_\** values.

The passive mechanical responses of the entire knee joint is governed by the mechanical properties of the four ligaments, namely the two collateral ligaments, MCL and LCL (medial and lateral collateral ligament), the cruciate ligaments, ACL and PCL (anterior and posterior cruciate ligaments), the patellar tendon, the medial and lateral menisci and articular cartilage [17]. The tendons and ligaments mainly act under tensile loads while the meniscus and cartilage support compression along with undergoing shear and transverse loads during movement. While there have been certain theoretical approaches like the lumped parameter models to simulate the knee joint as an ensemble of the underlying tissues, these models face certain restrictions, in incorporating the tissue geometry and their nonlinear elastic properties [18]. Further the current multiscale computational models are based on tissue properties sampled from different studies comprising of different animal models and individuals, which reduce their reliability due to variability in sampling, specimen size and condition, and variation in measurement techniques [19, 20]. However, to our current knowledge, no studies adopt a multiscale, experimental approach to evaluate the knee-joint biomechanics by determining the individual structural and functional properties of the underlying soft tissues and laxity properties of the same joint in ovine models.

For a deeper understanding of some of these open questions, in this study we investigate the relationships between whole joint and individual tissue properties using a combination of mechanical testing and imaging techniques. Specifically, we compare the results from joint laxity measurements, parameters from joint morphology, individual tissue structure and mechanical properties, collected from the anterior and posterior cruciate ligaments (ACL, PCL), medial and lateral collateral ligaments (MCL, LCL), the patellar tendon, menisci, and femoral cartilage from six different ovine specimens. We combine mechanical testing at two length scales with anatomical and quantitative MRI scans, to investigate whether there are any significant correlations between the individual tissue and the whole joint mechanical properties. Together, our results reveal relationships between whole joint mechanics, joint size, and individual tissue properties. We show positive correlations between the joint laxity forces and the inter-epicondylar distance for the tested ovine specimens and observe direct correlations with the ligament and tendon tissue mechanics. These results can provide insights into the differential role of individual tissue properties to the overall joint responses that will help design sample-specific multiscale models to simulate the knee joint biomechanics more accurately. The framework of these studies can be extended in future to human tissues to help design better strategies for improved joint health from a clinical perspective.

## METHODS

### Ovine stifle collection and storage

Ovine specimens were selected for this study owing to their physiological similarities with humans such as comparable body mass, similarities in neuroanatomical structures to humans and equivalent distribution of mechanical loads acting across the joints during load-bearing activities that resembles humans [6, 21]. Further the relative low-cost in comparison to other large animals, use of similar orthopedic instrumentation to humans for surgical approaches and having a comparable bone healing rate to that of humans makes them a useful large animal model in the field of musculoskeletal diseases. [6, 22]. Ovine stifles were obtained after humane euthanasia of sheep used for unrelated teaching purposes under institutional approval (IACUC protocol 1112000352) before preparation for multi-scale experiments (Figure 1). Skin and most of the musculature were removed except for those connected to the quadriceps and patellar tendon, to keep the joint capsule intact and not compromise with the joint stability. The distal femur and proximal tibia were cut mid-diaphysis and cleaned for ease of potting during whole joint experiments. The collected stifles were cleaned, wrapped in cotton gauze soaked in phosphate-buffered saline (PBS) to maintain hydration, and stored in plastic bags at −20°C. The frozen ovine stifles were thawed at 4°C for 24-36 hours before mechanical testing and MRI.

**Figure 1.**
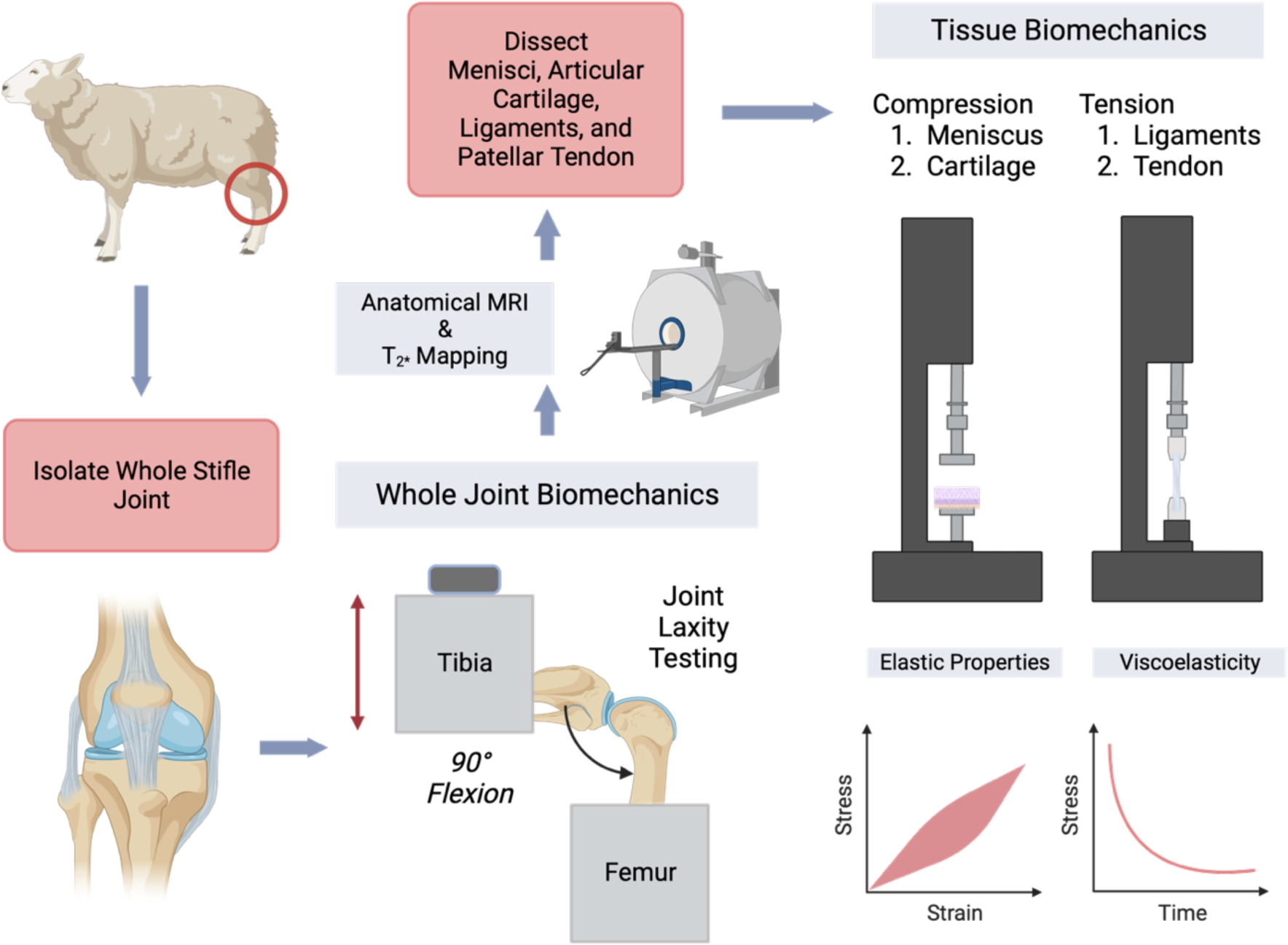
A schematic showing the experimental workflow for multiscale biomechanical testing and imaging of post-mortem ovine stifles. Post-collection of the ovine stifles, the excess musculature was removed, keeping the joint capsule intact and the samples were prepared for Joint-Laxity Testing. Following laxity testing, the stifles were imaged in MRI to quantify joint anatomy and *T_2_\** relaxation maps for tissues including the cartilage, menisci, ligaments, and patellar tendon. After imaging, the stifles were dissected, and the individual tissues were extracted in specific geometries suitable for mechanical testing. Cartilage and Menisci samples were tested under compression while tendons and ligaments were tested under tensile loading. The corresponding data was used to quantify the elastic and viscoelastic properties of the tissue explants. Schematic created with Biorender.com.

### Joint laxity testing

Joint laxity testing (**Figure 1**) was performed on ovine stifles using a 10-kN capacity load frame (MTS Systems). Prior to mechanical testing, the distal end of the femur and the proximal end of the tibia were cleared of tissue and embedded in cylindrical aluminum tubes using polymethyl methacrylate (Coralite Dental Manufacturing), mixed at 3:1 powder-to-liquid ratio and cured for 30 minutes. The stifle was wrapped in PBS-soaked gauze to prevent dehydration of the tissues during the curing time. The potted joints were attached to custom-built testing fixtures (**Figure 2a**) and mounted on the load frame at a 90° flexion angle [23]. Each stifle was preconditioned for twenty cycles using displacement control mode, and sinusoidal displacements of ±0.5 mm was provided at a rate of 0.2mm/s [9]. Following preconditioning, the joint was allowed to rest in its neutral position for two minutes and then loaded cyclically to ±1.5-mm sinusoidal displacements for 10 cycles at 0.2mm/s (**Figure 2b**). The difference between the peak forces at the endpoints of the cyclic displacement input was calculated as the range of forces experienced during testing and was used as a measure of joint laxity.

**Figure 2.**
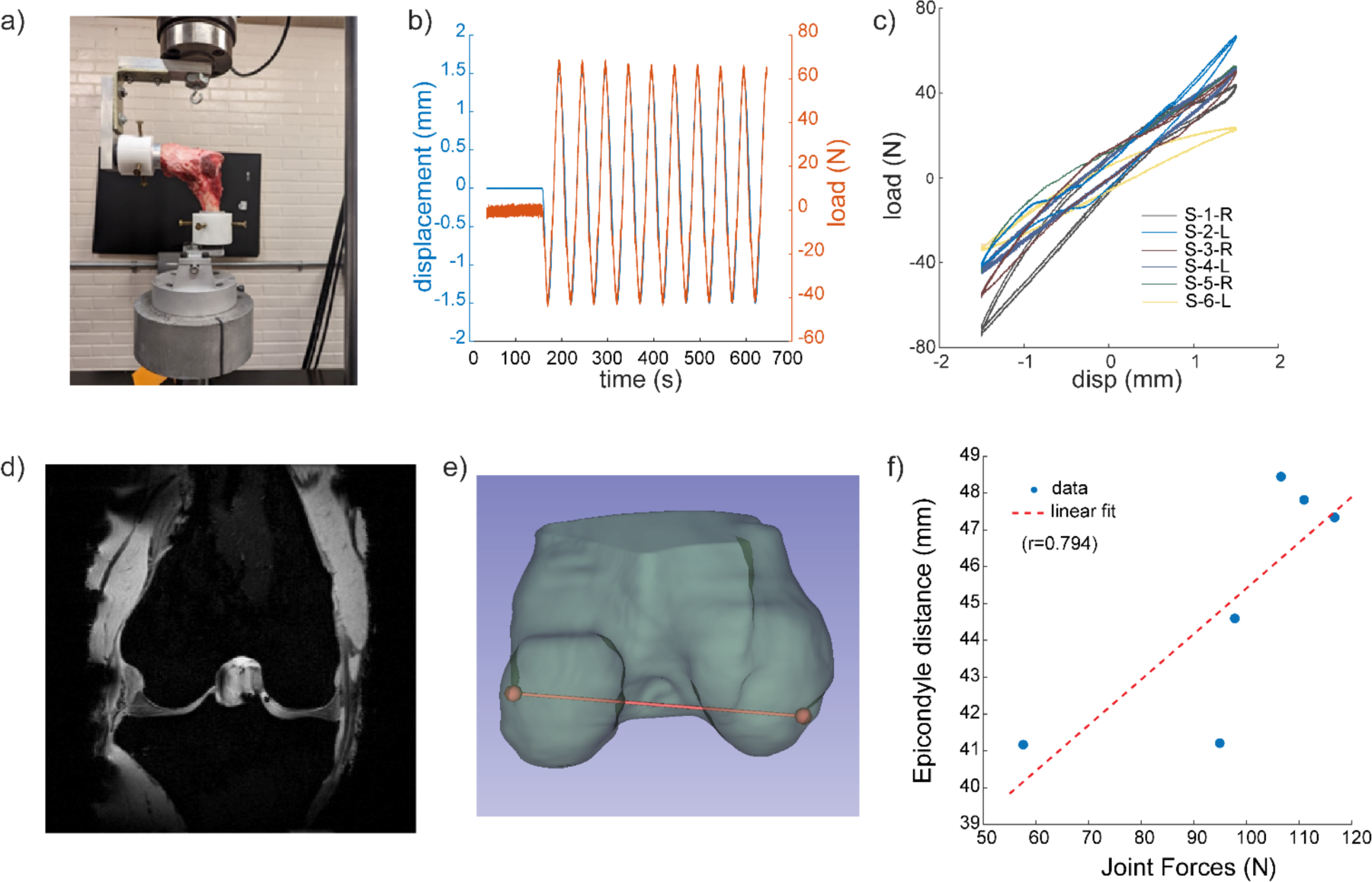
Measurement of Joint Laxity forces and its correlations with stifle size. A) Ovine stifles were clamped at 90° flexion to a 10-kN load frame for joint laxity testing. B) Cyclic load (N) and displacement (mm) profiles over time obtained during laxity test. C) Corresponding load vs displacements relationships were quantified during joint laxity tests for 6 ovine stifles. D) 3D *T_1_*-weighted FLASH images (representative image from specimen S1-R) were used to segment the femur. E) Epicondyle-to-epicondyle distance was measured from the segmented femur. F) Joint laxity forces positively correlated with epicondylar distance.

### MRI for Anatomy and T_2_* Mapping

The ovine stifles were imaged (**Figure 1**) using a Bruker Biospec 70/30 7T MRI (Billerica, MA) running manufacturer’s software (ParaVision 6.0.1). Each stifle was wrapped in gauze soaked in 1x PBS gauze to keep a hydrated environment during scanning. 3D *T_1_*-weighted FLASH scans [8-ms echo time (TE), 50-ms repetition time (TR), 20° flip angle, voxel size = 1/3 mm isotropic] were acquired to visualize tissue geometry (**Figure 2d**). A 2D multi-slice multi-echo gradient echo sequence was used to determine the *T_2_\** relaxation value of the tissues [TEs at 3.5 to 58.5 ms, at 5 ms echo spacing, TR = 1500 ms, in-plane spatial resolution = 1/3 mm × 1/3 mm, flip angle = 50°, slice thickness = 1 mm]. Joint anatomy was manually segmented in 3D Slicer (Version 5.2.2) software [24], and the femoral epicondyle-to-epicondyle distance was determined in Slicer (**Figure 2e**). The *T_2_\** analysis was performed using all the acquired slices from the 3-D scans of individual tissues for each medial meniscus, lateral meniscus, articular cartilage from the medial and lateral femoral condyles, the anterior and posterior cruciate ligaments (ACL, PCL), medial and lateral collateral ligaments (MCL, LCL), and the patellar tendon. The signal intensity from the multi-TE images were fit to equation *S* = 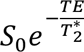 to estimate *T_2_\** relaxation times in MATLAB (Version 2022B. Natick, MA). Reported *T_2_\** averages include all pixels that had an exponential fit with an R^2^ value above 0.7.

### Tissue-specific mechanical testing to quantify material properties

Following the whole joint testing and MRI, each stifle was dissected to extract individual tissues for mechanical testing (**Figure 1**). Tissue specimens of standardized geometries suitable for mechanical testing were dissected from the ligaments (ACL, PCL, MCL, LCL) the patellar tendon, menisci, and articular cartilage from the femoral condyles of the tested ovine stifles. Prior to mechanical testing, tissue explants were equilibrated in PBS supplemented with 10 µL/mL protease inhibitor cocktail (Halt^TM^, Thermo Fisher Scientific) and 10 µL/mL 0.5-M EDTA (Thermo Fisher Scientific) at 4°C overnight.

For the cartilage and meniscus, cylindrical shaped explants were obtained using a bone punch (6-mm diameter) and a razor blade to trim to even thickness before compression testing [25, 26]. Cartilage and menisci explants were adhered to a 35-mm petri dish using cyanoacrylate and submerged in a PBS bath during testing. After preconditioning of 20 cycles to 5% compressive strain, at a rate of 1%/s of the gauge length [26], stress relaxation experiments were performed on the tissue specimens at 5 and 10% compressive strain to quantify viscoelastic behavior. The dwell time was kept at 30 minutes at each strain step for the explants to reach a steady state [27].

Tendon and ligament specimens were dissected to a target gauge length of ∼10 mm, width of ∼5 mm, and thickness of ∼ 2mm [28]. Preconditioning of 20 cycles to 5% tensile strain at 0.05 mm/s [29] was conducted. Stress relaxation tests were then performed to 5 and 10% tensile strain, with a dwell time of 20 minutes, [30] to quantify their viscoelastic properties.

### Estimation of tissue viscoelastic parameters from experimental data

The load vs time data obtained from the stress relaxation tests was converted to the corresponding stress vs time data using the information about sample geometry for each specimen. These data sets were used to estimate the viscoelastic properties of the explants using a 3-parameter nonlinear Prony Series model [31]:

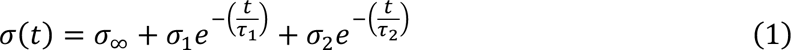

*σ*_∞_, *σ*_1_, and *σ*_2_ are stress parameters and *τ*_1_ and *τ*_2_ are their respective relaxation time constants. Parameters from the Prony series model were used to quantify the instantaneous modulus (*E*_0_)

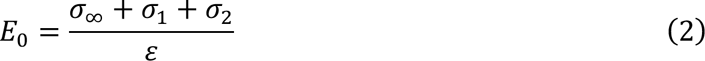

and relaxation modulus (*E*_∞_)

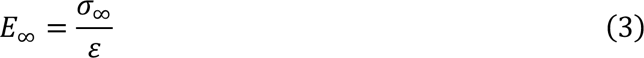

where *ε* is the applied strain [32]. In our analysis, we have defined the shorter time constant among *τ*_1_ and *τ*_2_ as *τ_min_* and the longer one as *τ_max_*. The Prony series model was fitted to experimental data using a non-linear least squares method in MATLAB.

### Statistical Analysis

All data are reported as mean ± standard deviation, unless otherwise indicated, and are publicly available for access [33]. Statistical analysis was performed using MATLAB, with a statistical significance defined at p < 0.05 for all hypothesis tests. The p-values were quantified using the *corrplot* function for two-tailed tests for the correlation coefficients. To quantify the strength of the correlations between different metrics, we quantified the Pearson’s linear correlation coefficient (r) between the joint laxity forces and the epicondyle-to-epicondyle distance, between normalized joint laxity forces and tissue moduli, and the *T_2_\** relaxation times. In our study, we have regarded the values of correlation coefficient (r), between 0.2-0.39 as weak, 0.40-0.59 as moderate, 0.6-0.79 as strong and 0.8-1 as significantly strong correlation.

## RESULTS

### Joint laxity measurements correlate with ovine stifle size

Laxity testing of the ovine stifles (**Figure 2**) under ten ±1.5-mm displacement cycles resulted in a 90.25 ± 32.33 N load, measured as the difference between the peak forces at the maximum and minimum displacements, across all six ovine specimens (**Figure 2c**). The femoral epicondyle-to-epicondyle distance positively correlated (r = 0.794, p = 0.059) to joint forces measured from the laxity experiments (**Figure 2f**). These results highlight the interdependencies between the knee joint size and its response to the external mechanical loading.

### T_2_*\** relaxation time does not significantly correlate to joint forces

*T_2_\** relaxation times were calculated for the stifle ligaments (ACL, PCL, MCL and LCL), cartilage, meniscus, and patellar tendon (**Figure 3**) and are reported as mean ± standard deviation of for each of the six ovine specimens (**Supplemental Table 1**). Within a particular tissue type, no statistically significant differences were observed in the mean *T_2_\** values among the six different specimens. Some moderate correlations were observed between the joint forces and tissue-specific *T_2_\** relaxation times for some tissues(**Figure 4a-b**, **Supplemental Table 2**): LCL (r=0.58, p=0.23), PCL (r=0.41,p=0.41), cartilage from the lateral femoral condyles (r=0.55, p=0.26), and lateral menisci (r=0.52, p=0.29); however, none of these reached statistical significance.

**Figure 3.**
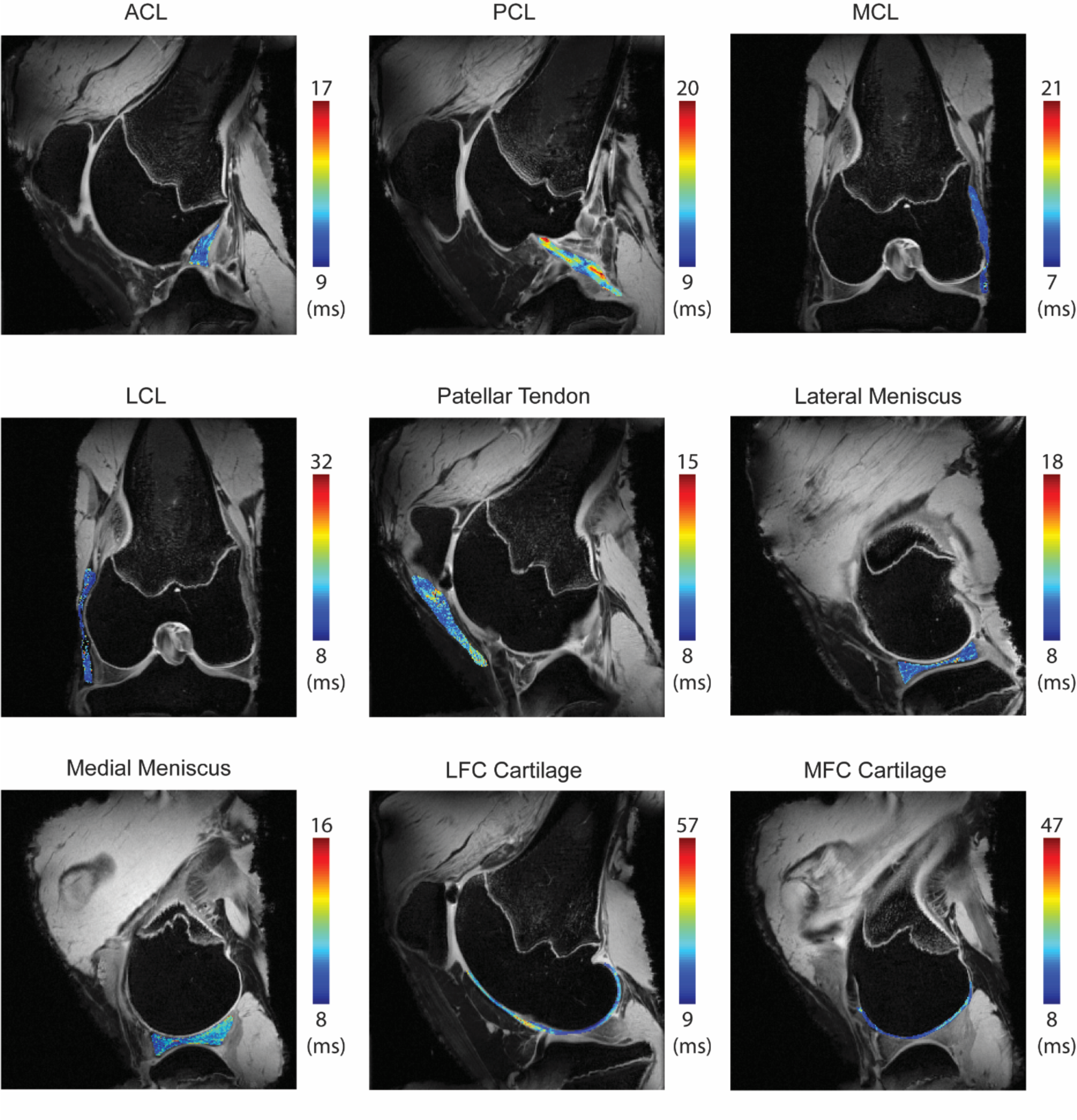
*T*_2_* maps were calculated from 2D multi-slice *T_2_\** multi gradient echo sequence and shown in representative slices for each tissue. Slices that show the *T_2_*\* map for the anterior cruciate ligament (ACL), posterior cruciate ligament (PCL), medial collateral ligament (MCL), lateral collateral ligament (LCL), patellar tendon, lateral and medial menisci, and cartilage from the lateral femoral condyle (LFC) and media femoral condyle (MFC) are shown for a representative stifle.

**Figure 4.**
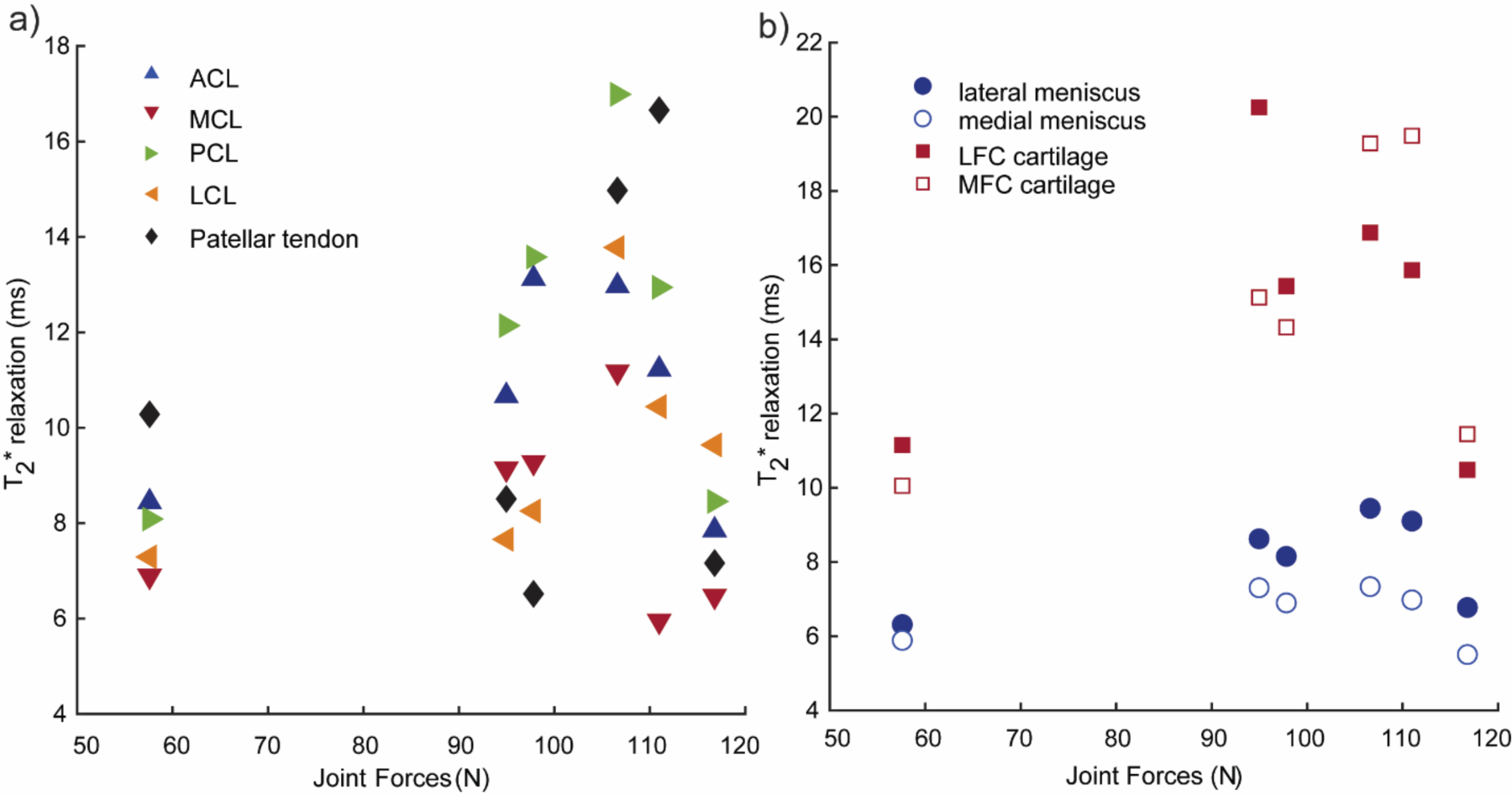
Joint forces (N)are compared with the measured mean *T_2_\** relaxation values (ms) for the different set of tissues. Joint forces and *T_2_\** relaxation time were evaluated for the patellar tendon and ligaments (ACL, PCL, MCL, LCL) (a) and for menisci and cartilage (b).

### Joint laxity forces positively correlate with tendon and ligament viscoelastic properties

We converted the load vs time data from the stress relaxation experiments on the patellar tendon and the four ligament tissues (ACL, PCL, MCL, LCL) under tensile loading to stress vs time data and used those to quantify instantaneous modulus (*E_o_*) and relaxation modulus (*E*_∞_) as measured under 5 and 10% strain (**Figure 5a-b**). The stress vs. time data is then fit to the 3-parameter Prony series model (R^2^ > 0.7) to obtain the tissue moduli and relaxation times (**Figure 5c**).

**Figure 5.**
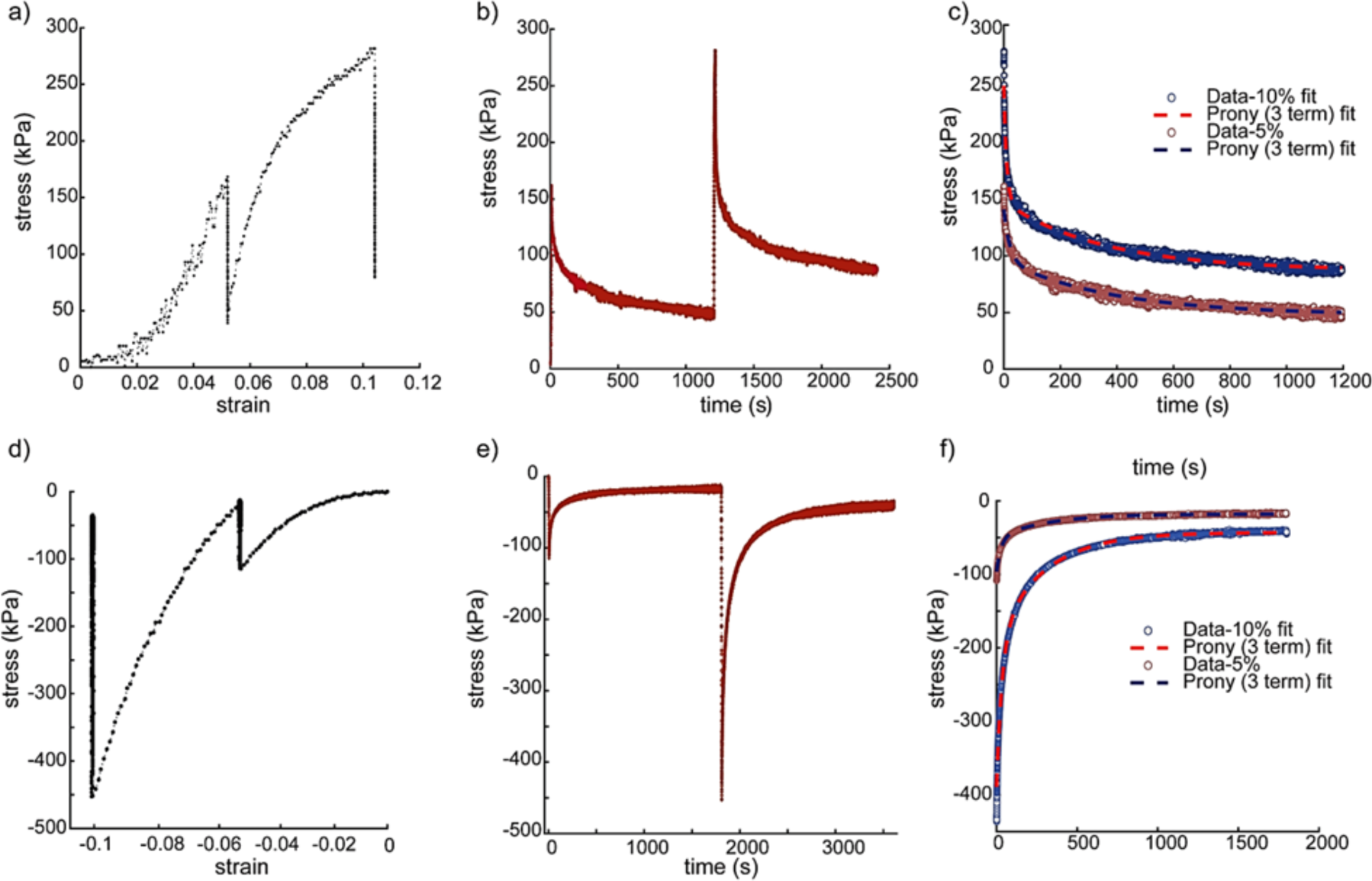
Representative stress-strain, stress-time profiles, and 3-parameter Prony series viscoelastic model fit to the experimental data under tensile and compressive loading. A lateral collateral ligament (LCL) sample was tested under 5 and 10% tensile strain (a-c). A medial meniscus sample was tested under 5 and 10% compressive strain (d-f).

We found strong positive correlations between joint laxity forces and both *E*_0_ and *E*_∞_of the tendons and ligaments (R^2^ > 0.7) but not cartilage or menisci (**Figure 6a-e**). Among the ligaments and the patellar tendon, the correlation coefficient values (r) between the tissue moduli and joint forces were comparatively stronger for the PCL and ACL. We also found statistically significant correlation among joint forces and viscoelastic parameters and weak to moderate correlations with T_2_*, specifically for the LCL, PCL, MCL and the patellar tendon; however, they were not statistically significant (**Figure 7a-e**, **Supplemental Table 2**). Overall, these results provide useful insights on how whole joint behavior may depend on individual properties of the tendons and ligaments present in the knee.

**Figure 6.**
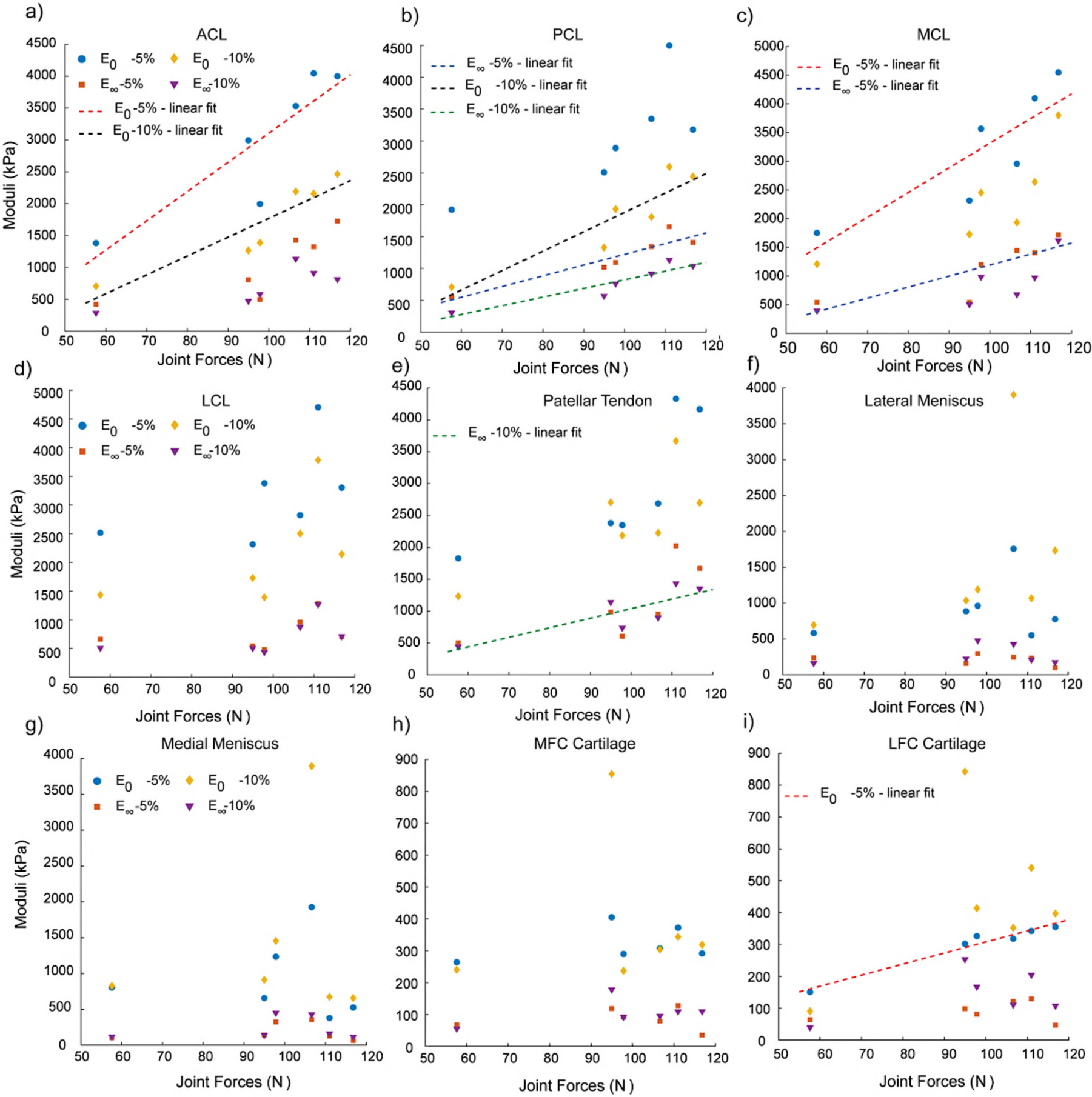
Variation in the tissue-specific viscoelastic properties (instantaneous and relaxation moduli (kPa) under 5 and 10% strain) for all the sets of tissues with the measured Joint laxity forces (N). Ligaments and tendons (a-e) show a positive trend with the laxity forces whereas f menisci and cartilage moduli (f-i) do not correlate with the joint forces. The linear fits denote strong positive correlations with Pearson’s linear correlation coefficient (R^2^ > 0.7).

**Figure 7.**
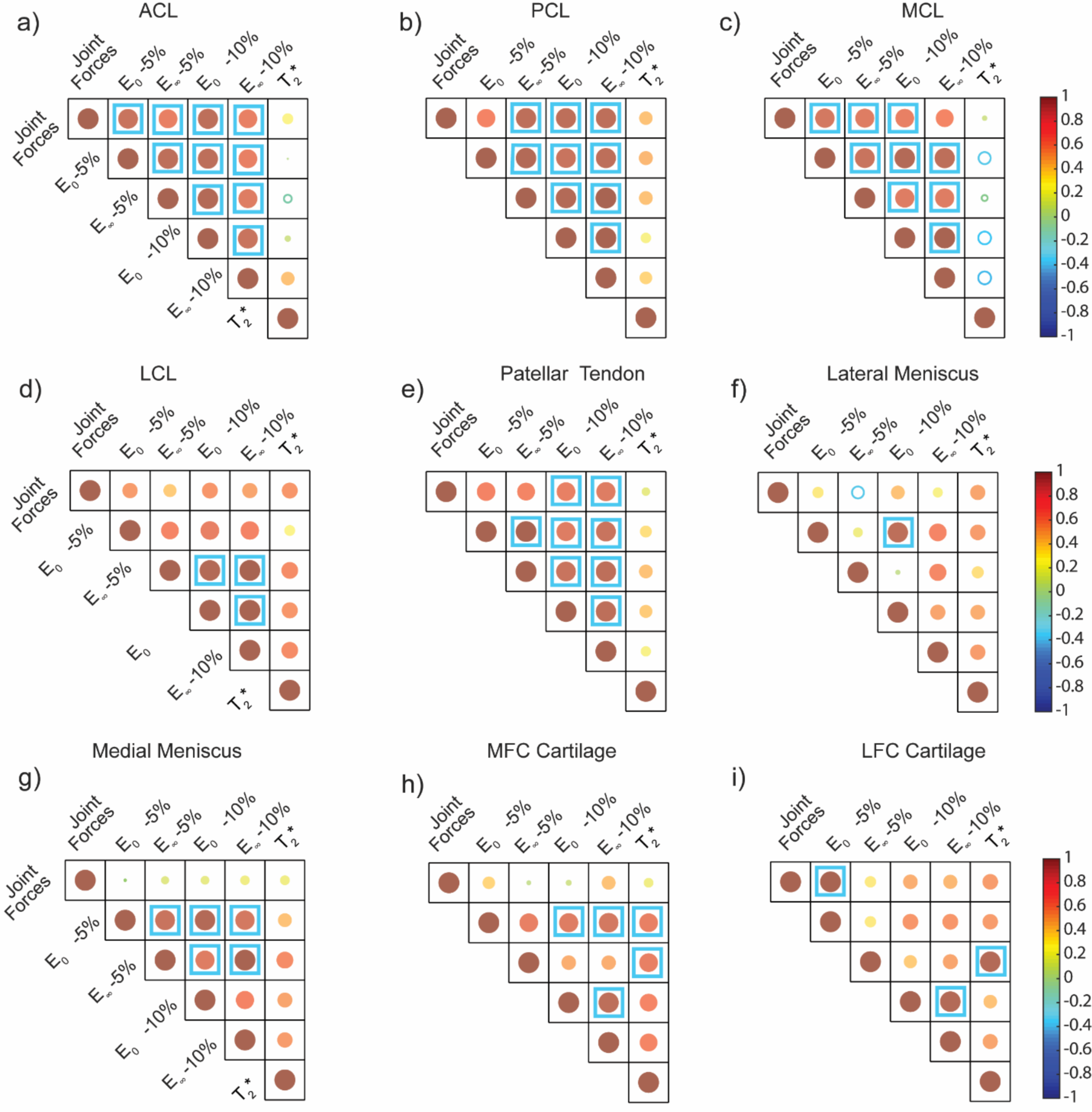
Correlation matrices plot for comparison among joint forces (N), viscoelastic properties, and *T_2_\** values for the different set of tested tissues. Viscoelastic properties – instantaneous and relaxation moduli (kPa) – were evaluated under 5% and 10% strain. *T_2_*\* relaxation times (ms) were measured for individual tissues using MRI. The size and color of the circles represent the Pearson’s linear correlation coefficient (r). Solid circles show a positive correlation (0 < r ≤1) and hollow circles show negative correlation (−1 ≤ r < 0). Any strongly positive correlations (r ≥ 0.8) are denoted with a blue square. The corresponding table of correlation coefficient (r) and p-values are provided in Supplemental Table 2.

### Joint Laxity measurements do not show a significant correlation with viscoelastic properties of cartilage and meniscus

Following a similar procedure as with the tendons and ligament data, we converted the load vs time data from the stress relaxation experiments on the femoral articular cartilage and menisci under compressive loading and used them to quantify the tissue *E_o_* and *E*_∞_under 5 & 10% strain using a 3-parameter Prony series model (**Figure 5d-f**). Unlike the tendons and ligaments, we did not observe any significant correlation with the whole-joint measurements and the cartilage and meniscus viscoelastic properties (**Figure 6f-i**), except for the *E_o_* at 5% strain of articular cartilage samples extracted from the lateral femoral condyle (r ≥ 0.95, p < 0.01). Articular cartilage extracted from both the lateral and medial femoral condyles showed strong positive correlation (r ≥ 0.8) between the instantaneous or relaxation moduli and *T_2_\** values (**Figure 7h-i**). Specifically, these correlations were observed between *E*_0_ at 10% and *T_2_\** (r =0.95, p = 0.003) and for both *E*_0_ and *E*_∞_ at 5% strain and *T_2_\** (both r = 0.8, p = 0.06). We also observed some moderately positive correlations between the tissue moduli measured at 10% strain and the *T_2_\** values for the lateral and medial menisci, although these correlations did not reach statistical significance (**Figure 7f-g**, **Supplemental Table 2**). These results highlight that the disparate response of different types of soft tissues within the knee may contribute to the overall joint response differently depending on the type of applied loading.

## DISCUSSION

The mechanical responses of the knee joint depend on multiple parameters, ranging from the structure of the underlying bones, the structure and mechanics of soft tissues that support the joint, applied load and motion constraints depending on the positioning of the joint [7]. Joint laxity, which inversely relate to joint stiffness, has previously been linked with the stiffness of individual internal structures using theoretical studies [34]. However, these theoretical studies often employing computational approaches like lumped parameter models [18] or finite element models [35], which require detailed inputs on the structural and mechanical properties of the underlying structures, often based on experimental data. Currently there remains a gap between the theoretical approaches and the available experimental data to determine these connections at multiple length scales. In this study, using a combination of multiscale biomechanical testing and MRI, we have evaluated the correlations between whole joint and tissue level mechanics with joint geometry and tissue *T_2_\** values that provide useful insights into the knee joint mechanics facilitating data interpretation for clinical evaluation.

In this study, we have conducted joint-laxity studies on ovine stifles, measuring the range of forces resulting under cyclic displacements, and used MRI to quantify the epicondylar distance as a proxy for joint size. We found a positive correlation between the epicondylar distance of the ovine stifles and the measured joint forces. In a previous study [36], statistical shape modelling showed that the morphological variabilities in the bone shape of tibia and femur and their relative alignment were linked with joint instability in human models. Such studies highlight the existence of underlying correlations between joint anatomy and biomechanics.

We also measured *T_2_\** of the different nonmineralized tissues of the stifle, including the ligaments, patellar tendon, menisci, and articular cartilage from the femoral condyles. The mean value of the *T_2_\** relaxation times measured for different tissue types and are in line with previously published studies. The average *T_2_\** values for the lateral and media menisci for all six specimens were 8.07± 2.93 ms and 6.65± 2.55 ms respectively which is similar to previous studies in literature [37]. The corresponding average *T_2_\** values of the cartilage from the lateral and medial femoral condyles measured in our study are 14.95 ± 7.17 ms and 15.01 ± 6.79 ms, which are lower than prior study showing *T_2_* of 30 ms [37]. We speculate that these variations can be due to the differences in magnetic field strengths of the MRI machines, in combination with variation in tissue properties dependent on various factors such the animal age, sex, body weight, and other parameters. Variations in *T_2_\** values could arise because of differences in positioning of the knee during scanning [38]. To reduce such variabilities in the measurements, we maintained a similar positioning of the knee joints for all the specimens during scanning. However, because the scans were performed in the intact joint, the differences in orientation of the primary direction of collagen fiber alignment in the tissues, with respect to the direction of the main magnetic field, cannot be fully controlled. The average *T_2_\** values of the patellar tendon and ligaments were 10.68 ± 4.48 (tendon), 10.71 ± 4.19 (ACL), 8.15 ± 3.55 (MCL), 12.03 ± 5.16 (PCL), 9.51 ± 4.92 (LCL) ms, in general agreement with an earlier study performed on tendon and ligaments from human cadaveric specimens [39]. In our study, we did not see any significant correlation between the tissue specific *T_2_\** values with the joint laxity forces; however, some moderately positive correlations were observed for few groups of ligaments, specifically the LCL and PCL, the cartilage from lateral femoral condyles and the lateral menisci. We hypothesize that adjusting these measurements with respect to the body weight may lead to improved correlation strengths, as reported in an earlier study [40]. These results corroborate with few earlier studies [41, 42] that have suggested possible correlations between the *T_2_\** relaxation time for specific tissues like cartilage and meniscus and joint health in diseased conditions such as osteoarthritis or ACL related injuries.

Finally, we quantified the mechanical properties of individual tissues and investigated their correlations to the whole joint laxity response. We found significantly positive correlations with the ligaments and patellar tendon moduli with the joint laxity forces. These results are in good agreement with previous studies [43, 44] that highlight that the knee joint responses are influenced by ligament mechanics. Further, in our study we observed the MCL to have the relatively highest instantaneous modulus among the different ligaments and tendons, that aligns with an earlier study [28] performed in bovine samples. While we do not see a significant correlation between the joint forces and the cartilage and meniscus mechanics, these observations may depend on the type of loading applied. Ligaments are crucial for knee joint stability as they provide mechanical reinforcements and help in motion control [45]. One limitation of our study is that we did not quantify the collagen and proteoglycan contents in the tested tissues. While there’s currently no clear consensus on the variation of net collagen or overall matrix content in tendons and knee joint ligaments [46, 47], variation in the water, proteoglycan and collagen content in bovine ligaments and tendons can be linked to changes in their mechanical properties [48, 49]. While we did not observe any significant correlations between the tissue moduli and the *T_2_\** for the tendon and ligaments, there were some moderately positive correlations was observed between *T_2_\** values and tissue moduli of the LCL and PCL, weak positive correlations for the patellar tendon and a weak inverse correlation for the MCL. These results are in line with a recently published study that observed significant correlations between the tissue modulus and *T_1ρ_* measured in Achilles tendons obtained from human cadaveric samples [50]. On the other hand, we observed that the articular cartilage extracted from the medial and femoral condyles of the ovine stifles exhibited a significantly positive correlation between the tissue mechanics and the *T_2_\** relaxation values. Viscoelastic properties of cartilage have been previously correlated with tissue degradation and early OA progression using canine cranial cruciate ligament transection models [51]; however, the correlations between the tissue viscoelastic properties with relaxometry methods such as *T_2_\** remain an area for further exploration in future [52].

In conclusion, we found correlations between whole joint mechanics, joint size, and individual tissue properties in ovine stifles. Specifically, forces measured during whole joint testing correlated directly with femur size, measured using the epicondylar distance. We also find that the viscoelastic properties of the tendons and ligaments correlated positively with joint laxity forces. We also observe a moderate inverse correlation with tissue viscoelastic properties with *T_2_\** for patellar tendon and positive correlations for cartilage samples from the femoral condyles. However, we did not find any other significant correlations between tissue *T_2_\** relaxation times and viscoelastic properties for the other groups of tissues. These results can provide useful insights into the differential role of individual tissue properties that can be used to design sample specific computational models to assess the knee joint properties as an ensemble of the underlying soft structures for development of better diagnostic techniques and clinical assessment.

## ACKNOWLEDGEMENTS

This work was funded in part by the Department of Defense (ARO W911NF2110372). The authors would also like to thank Ms. Shreya Sinha for help in sample preparation for the joint-laxity tests.

