## Supplemental Table 1 for "Multiscale Correlations between Joint and Tissue-Specific Biomechanics and Anatomy in Postmortem Ovine Stifles"

**Supplemental Table 1. *T_2_^*^* relaxation values in ms (mean ± std) for the menisci, cartilage (from either medial and lateral femoral condyles), ligaments and tendon for the tested ovine specimens (total six – one stifle per each animal)**

| **Specimen** | **Lateral Meniscus** | **Medial Meniscus** | **Cartilage (LFC)** | **Cartilage (MFC)** | **ACL** | **MCL** | **PCL** | **LCL** | **Patellar Tendon** |
| --- | --- | --- | --- | --- | --- | --- | --- | --- | --- |
| **S-1** | 6.77 ±2.66 | 5.51±1.81 | 11.45±6.4 | 10.48±5.61 | 7.85±2.73 | 6.47±2.11 | 8.46±4.19 | 9.64±5.15 | 7.16±3.27 |
| **S-2** | 9.1±3.71 | 6.98±2.76 | 19.49±10.5 | 15.87±6.25 | 11.22±4.35 | 5.95±2.28 | 12.95±6.08 | 10.44±6.07 | 16.66±6.92 |
| **S-3** | 9.45±3.81 | 7.34±3.39 | 19.28±9.14 | 16.88±6.46 | 12.97±5.58 | 11.17±6.95 | 16.99±6.79 | 13.78±8.91 | 14.97±6.17 |
| **S-4** | 8.15±2.36 | 6.9±2.46 | 14.33±4.04 | 15.43±6.55 | 13.13±4.58 | 9.28±3.38 | 13.58±6.1 | 8.26±3.11 | 6.52±2.76 |
| **S-5** | 8.63±2.87 | 7.31±2.07 | 15.13±6.69 | 20.25±8.5 | 10.67±3.81 | 9.14±3.81 | 12.14±3.67 | 7.66±2.21 | 8.51±2.59 |
| **S-6** | 6.32±2.16 | 5.89±2.82 | 10.06±6.22 | 11.15±7.4 | 8.44±4.06 | 6.89±2.74 | 8.09±4.09 | 7.29±4.09 | 10.28±5.17 |

**Supplemental Table 2. Tissue-specific correlation coefficient (r) and p-values calculated from comparison between joint forces (N), viscoelastic properties (instantaneous (**$E_{o}$**) and relaxation (**$E_{\infty}$**) moduli (kPa) under 5 & 10% strain), and *T_2_^*^* relaxation times (ms).** The upper diagonal terms represent the correlation coefficient (r), and the lower diagonal terms represent the corresponding p-values (in italics). Significant p-values and their corresponding correlation coefficients are underlined.

| **Anterior Cruciate Ligament** | | | | | | |
| --- | --- | --- | --- | --- | --- | --- |
| **r**  **p** | **Joint Forces (N)** | $\boldsymbol{E}_{\boldsymbol{o}}$**, -5%** | $\boldsymbol{E}_{\boldsymbol{\infty}}$**, -5%** | $\boldsymbol{E}_{\boldsymbol{o}}$**, -10%** | $\boldsymbol{E}_{\boldsymbol{\infty}}$**, -10%** | **Average *T_2_^*^* (ms)** |
| **Joint Forces (N)** | 1 | 0.88 | 0.81 | 0.92 | 0.80 | 0.27 |
| $\boldsymbol{E}_{\boldsymbol{o}}$**, -5%** | *0.019* | 1 | 0.93 | 0.92 | 0.80 | 0.01 |
| $\boldsymbol{E}_{\boldsymbol{\infty}}$**, -5%** | *0.052* | *0.006* | 1 | 0.95 | 0.82 | -0.14 |
| $\boldsymbol{E}_{\boldsymbol{o}}$**, -10%** | *0.009* | *0.009* | *0.003* | 1 | 0.89 | 0.09 |
| $\boldsymbol{E}_{\boldsymbol{\infty}}$**, -10%** | *0.060* | *0.058* | *0.047* | *0.016* | 1 | 0.42 |
| **Average *T_2_^*^* (ms)** | *0.601* | *0.984* | *0.792* | *0.865* | *0.414* | 1 |
| **Posterior Cruciate Ligament** | | | | | | |
| **r**  **p** | **Joint Forces (N)** | $\boldsymbol{E}_{\boldsymbol{o}}$**, -5%** | $\boldsymbol{E}_{\boldsymbol{\infty}}$**, -5%** | $\boldsymbol{E}_{\boldsymbol{o}}$**, -10%** | $\boldsymbol{E}_{\boldsymbol{\infty}}$**, -10%** | **Average *T_2_^*^* (ms)** |
| **Joint Forces (N)** | 1 | 0.77 | 0.93 | 0.91 | 0.93 | 0.41 |
| $\boldsymbol{E}_{\boldsymbol{o}}$**, -5%** | *0.072* | 1 | 0.95 | 0.88 | 0.92 | 0.44 |
| $\boldsymbol{E}_{\boldsymbol{\infty}}$**, -5%** | *0.007* | *0.004* | 1 | 0.95 | 0.98 | 0.43 |
| $\boldsymbol{E}_{\boldsymbol{o}}$**, -10%** | *0.011* | *0.020* | *0.004* | 1 | 0.97 | 0.25 |
| $\boldsymbol{E}_{\boldsymbol{\infty}}$**, -10%** | *0.007* | *0.01* | *0.0004* | *0.001* | 1 | 0.37 |
| **Average *T_2_^*^* (ms)** | *0.417* | *0.377* | *0.396* | *0.635* | *0.476* | 1 |
| **Lateral Collateral Ligament** | | | | | | |
| **r**  **p** | **Joint Forces (N)** | $\boldsymbol{E}_{\boldsymbol{o}}$**, -5%** | $\boldsymbol{E}_{\boldsymbol{\infty}}$**, -5%** | $\boldsymbol{E}_{\boldsymbol{o}}$**, -10%** | $\boldsymbol{E}_{\boldsymbol{\infty}}$**, -10%** | **Average *T_2_^*^* (ms)** |
| **Joint Forces (N)** | 1 | 0.78 | 0.72 | 0.80 | 0.83 | 0.58 |
| $\boldsymbol{E}_{\boldsymbol{o}}$**, -5%** | *0.067* | 1 | 0.97 | 0.82 | 0.89 | 0.26 |
| $\boldsymbol{E}_{\boldsymbol{\infty}}$**, -5%** | *0.103* | *0.001* | 1 | 0.88 | 0.93 | 0.64 |
| $\boldsymbol{E}_{\boldsymbol{o}}$**, -10%** | *0.061* | *0.045* | *0.020* | 1 | 0.93 | 0.59 |
| $\boldsymbol{E}_{\boldsymbol{\infty}}$**, -10%** | *0.039* | *0.017* | *0.007* | *0.006* | 1 | 0.63 |
| **Average *T_2_^*^* (ms)** | *0.231* | *0.613* | *0.174* | *0.222* | *0.179* | 1 |
| **Medial Collateral Ligament** | | | | | | |
| **r**  **p** | **Joint Forces (N)** | $\boldsymbol{E}_{\boldsymbol{o}}$**, -5%** | $\boldsymbol{E}_{\boldsymbol{\infty}}$**, -5%** | $\boldsymbol{E}_{\boldsymbol{o}}$**, -10%** | $\boldsymbol{E}_{\boldsymbol{\infty}}$**, -10%** | **Average *T_2_^*^* (ms)** |
| **Joint Forces (N)** | 1 | 0.85 | 0.82 | 0.79 | 0.73 | 0.06 |
| $\boldsymbol{E}_{\boldsymbol{o}}$**, -5%** | *0.032* | 1 | 0.90 | 0.95 | 0.93 | -0.33 |
| $\boldsymbol{E}_{\boldsymbol{\infty}}$**, -5%** | *0.044* | *0.016* | 1 | 0.83 | 0.83 | -0.08 |
| $\boldsymbol{E}_{\boldsymbol{o}}$**, -10%** | *0.061* | *0.004* | *0.039* | 1 | 0.99 | -0.37 |
| $\boldsymbol{E}_{\boldsymbol{\infty}}$**, -10%** | *0.102* | *0.007* | *0.039* | *0.0001* | *1* | -0.38 |
| **Average *T_2_^*^* (ms)** | *0.91* | *0.518* | *0.879* | *0.468* | *0.462* | 1 |
| **Patellar Tendon** | | | | | | |
| **r**  **p** | **Joint Forces (N)** | $\boldsymbol{E}_{\boldsymbol{o}}$**, -5%** | $\boldsymbol{E}_{\boldsymbol{\infty}}$**, -5%** | $\boldsymbol{E}_{\boldsymbol{o}}$**, -10%** | $\boldsymbol{E}_{\boldsymbol{\infty}}$**, -10%** | **Average *T_2_^*^* (ms)** |
| **Joint Forces (N)** | 1 | 0.78 | 0.723 | 0.8 | 0.83 | 0.16 |
| $\boldsymbol{E}_{\boldsymbol{o}}$**, -5%** | *0.067* | 1 | 0.97 | 0.82 | 0.89 | 0.32 |
| $\boldsymbol{E}_{\boldsymbol{\infty}}$**, -5%** | *0.103* | *0.001* | 1 | 0.88 | 0.93 | 0.43 |
| $\boldsymbol{E}_{\boldsymbol{o}}$**, -10%** | *0.061* | *0.045* | *0.020* | 1 | 0.93 | 0.38 |
| $\boldsymbol{E}_{\boldsymbol{\infty}}$**, -10%** | *0.039* | *0.017* | *0.007* | *0.006* | 1 | 0.24 |
| **Average *T_2_^*^* (ms)** | *0.76* | *0.534* | *0.397* | *0.453* | *0.648* | 1 |
| **Lateral Femoral Condylar Cartilage** | | | | | | |
| **r**  **p** | **Joint Forces (N)** | $\boldsymbol{E}_{\boldsymbol{o}}$**, -5%** | $\boldsymbol{E}_{\boldsymbol{\infty}}$**, -5%** | $\boldsymbol{E}_{\boldsymbol{o}}$**, -10%** | $\boldsymbol{E}_{\boldsymbol{\infty}}$**, -10%** | **Average *T_2_^*^* (ms)** |
| **Joint Forces (N)** | 1 | 0.98 | 0.29 | 0.48 | 0.46 | 0.55 |
| $\boldsymbol{E}_{\boldsymbol{o}}$**, -5%** | *0.0007* | 1 | 0.29 | 0.57 | 0.57 | 0.54 |
| $\boldsymbol{E}_{\boldsymbol{\infty}}$**, -5%** | *0.573* | *0.575* | 1 | 0.39 | 0.51 | 0.95 |
| $\boldsymbol{E}_{\boldsymbol{o}}$**, -10%** | *0.330* | *0.240* | *0.439* | 1 | 0.96 | 0.42 |
| $\boldsymbol{E}_{\boldsymbol{\infty}}$**, -10%** | *0.363* | *0.239* | *0.299* | *0.003* | 1 | 0.52 |
| **Average *T_2_^*^* (ms)** | *0.256* | *0.267* | *0.003* | *0.406* | *0.292* | 1 |
| **Medial Femoral Condylar Cartilage** | | | | | | |
| **r**  **p** | **Joint Forces (N)** | $\boldsymbol{E}_{\boldsymbol{o}}$**, -5%** | $\boldsymbol{E}_{\boldsymbol{\infty}}$**, -5%** | $\boldsymbol{E}_{\boldsymbol{o}}$**, -10%** | $\boldsymbol{E}_{\boldsymbol{\infty}}$**, -10%** | **Average *T_2_^*^* (ms)** |
| **Joint Forces (N)** | 1 | 0.35 | 0.05 | 0.08 | 0.44 | 0.211 |
| $\boldsymbol{E}_{\boldsymbol{o}}$**, -5%** | *0.491* | 1 | 0.79 | 0.83 | 0.87 | 0.80 |
| $\boldsymbol{E}_{\boldsymbol{\infty}}$**, -5%** | *0.922* | *0.063* | 1 | 0.48 | 0.47 | 0.80 |
| $\boldsymbol{E}_{\boldsymbol{o}}$**, -10%** | *0.882* | *0.041* | *0.334* | 1 | 0.93 | 0.71 |
| $\boldsymbol{E}_{\boldsymbol{\infty}}$**, -10%** | *0.387* | *0.024* | *0.346* | *0.008* | 1 | 0.72 |
| **Average *T_2_^*^* (ms)** | *0.687* | *0.059* | *0.059* | *0.112* | *0.104* | 1 |
| **Lateral Meniscus** | | | | | | |
| **r**  **p** | **Joint Forces (N)** | $\boldsymbol{E}_{\boldsymbol{o}}$**, -5%** | $\boldsymbol{E}_{\boldsymbol{\infty}}$**, -5%** | $\boldsymbol{E}_{\boldsymbol{o}}$**, -10%** | $\boldsymbol{E}_{\boldsymbol{\infty}}$**, -10%** | **Average *T_2_^*^* (ms)** |
| **Joint Forces (N)** | 1 | 0.22 | -0.40 | 0.29 | 0.20 | 0.52 |
| $\boldsymbol{E}_{\boldsymbol{o}}$**, -5%** | *0.677* | 1 | 0.22 | 0.93 | 0.71 | 0.55 |
| $\boldsymbol{E}_{\boldsymbol{\infty}}$**, -5%** | *0.430* | *0.677* | 1 | 0.06 | 0.67 | 0.33 |
| $\boldsymbol{E}_{\boldsymbol{o}}$**, -10%** | *0.580* | *0.007* | *0.914* | 1 | 0.52 | 0.50 |
| $\boldsymbol{E}_{\boldsymbol{\infty}}$**, -10%** | *0.705* | *0.114* | *0.148* | *0.288* | 1 | 0.54 |
| **Average *T_2_^*^* (ms)** | *0.287* | *0.260* | *0.524* | *0.318* | *0.273* | 1 |
| **Medial Meniscus** | | | | | | |
| **r**  **p** | **Joint Forces (N)** | $\boldsymbol{E}_{\boldsymbol{o}}$**, -5%** | $\boldsymbol{E}_{\boldsymbol{\infty}}$**, -5%** | $\boldsymbol{E}_{\boldsymbol{o}}$**, -10%** | $\boldsymbol{E}_{\boldsymbol{\infty}}$**, -10%** | **Average *T_2_^*^* (ms)** |
| **Joint Forces (N)** | 1 | -0.01 | 0.15 | 0.17 | 0.21 | 0.21 |
| $\boldsymbol{E}_{\boldsymbol{o}}$**, -5%** | *0.99* | 1 | 0.90 | 0.95 | 0.85 | 0.43 |
| $\boldsymbol{E}_{\boldsymbol{\infty}}$**, -5%** | *0.781* | *0.015* | 1 | 0.83 | 0.98 | 0.65 |
| $\boldsymbol{E}_{\boldsymbol{o}}$**, -10%** | *0.745* | *0.004* | *0.043* | 1 | 0.75 | 0.51 |
| $\boldsymbol{E}_{\boldsymbol{\infty}}$**, -10%** | *0.695* | *0.031* | *0.0005* | *0.085* | 1 | 0.53 |
| **Average *T_2_^*^* (ms)** | *0.683* | *0.39* | *0.166* | *0.303* | *0.274* | 1 |
